# Molecular signature of RV cardiac remodeling associated with Polymerase Gamma mutation

**DOI:** 10.1101/2022.02.11.480145

**Authors:** Matthew W. Gorr, Ashley Francois, Lynn M. Marcho, Ty Saldana, Erin McGrail, Nuo Sun, Matthew S. Stratton

## Abstract

Aging is associated with metabolic dysfunction and metabolic dysfunction accelerates the course of aging. This is well illustrated by aging-like phenotypes displayed in the Polymerase Gamma mutant mouse. Here, a key residue of the mitochondrial DNA polymerase is mutated (D257A) which hinders proof reading capacity, resulting in mitochondrial DNA mutation and instability. Given known cardiac phenotypes in the POLG mutant mouse, we sought to characterize the course of cardiac dysfunction in the POLG mutant to guide future intervention studies. Including both male and female animals allowed interrogation of sex-specific responses to severe metabolic dysfunction. We also conducted RNA-seq analysis on cardiac right ventricles to broadly identify mechanisms engaged by severe metabolic dysfunction that are associated with RV pathology. Several interesting sex differences were noted. Specifically, female POLG mutants died earlier than male POLG mutants and LV chamber diameters were impacted earlier in females than males. Moreover, male POLG mutants showed LV wall thinning while female POLG mutant LV walls were thicker. Both males and females displayed significant RV hypertrophy. Finally, RNA-seq analysis of the RV tissue indicated that POLG mutation significantly impacted canonical pathways associated with inflammation, fibrosis, and heart failure. Comparison of this RNA-seq dataset with other publically available datasets highlight 1) strong conservation among downregulated genes in models of accelerated cardiac aging, 2) the potential involvement of the mitochondrial unfolded protein response in the POLG mutant, and 3) the ability of zinc dependent HDAC inhibition to rescue the expression of genes downregulated in accelerated cardiac aging.

## Introduction

Aging is a complex process that impacts all organ systems of the body, and their interactions. Many cellular and molecular mechanisms have been implicated in the biology of aging, including chronic low-grade inflammation, cellular senescence, DNA damage, telomere shortening, mTOR signaling, ROS generation, loss of proteostasis, and mitochondrial dysfunction (<u>1</u>). These mechanisms are related and subject to significant environmental influence (<u>2</u>). Mitochondrial function and mitochondria-derived signals are central to aging pathology (<u>3</u>, <u>4</u>). With an eye toward feasibility of cardiac aging intervention studies in a pandemic environment, we sought a model of accelerated aging given advantages in defining shorter intervention/treatment time windows and robust phenotypic changes to facilitate unbiased approaches to identify aging-associated mechanisms.

Mutations in the mitochondrial DNA polymerase, Polymerase Gamma (POLG) have been associated with a number of mitochondrial diseases in humans (reviewed in <u>5</u>). Several POLG mutant mouse models have been developed that broadly display accelerated aging phenotypes and cardiac remodeling consistent with accelerated cardiac aging (e.g. <u>6–9</u>). The POLG D257A point mutant mouse lacks proof reading capacity in the mitochondrial DNA polymerase, resulting in reduced mitochondrial DNA integrity (<u>10</u>). These mice have been reported to display phenotypes consistent with accelerated cardiac aging, though literature reporting the degree of cardiac dysfunction in these animals is varied (6, 11-14).

## Results

As previously reported, POLG mutants displayed gross phenotypes consistent with accelerated-aging, including hair loss, weight loss, frail bones, kyphosis, and early mortality (6). Given know variations in disease pathology/progression based on environmental factors that vary from institution to institution (e.g. diet, light cycle, housing conditions and density, vivarium traffic, elevation, etc.) we embarked on a pilot survival study to determine the POLG life-span locally. Here, we include both natural death and veterinarian directed euthanasia due to severely deteriorated condition as death (lack of response to stimulation, loss of >20% of body weight, Body Condition Score <2). The average age of death for male POLG mice was 322.5 days of age (+/−8.154, N=14) while female average survival was only to 276.7 days (Figure 1A). These data are in agreement with significantly worsened outcomes in a classical survival analysis in the female relative to male POLG mutants (Figure 1B, Log-rank test p=0.0026 and Gehan-Breslow-Wokcoxon test p=0.0073).

**Figure 1.**
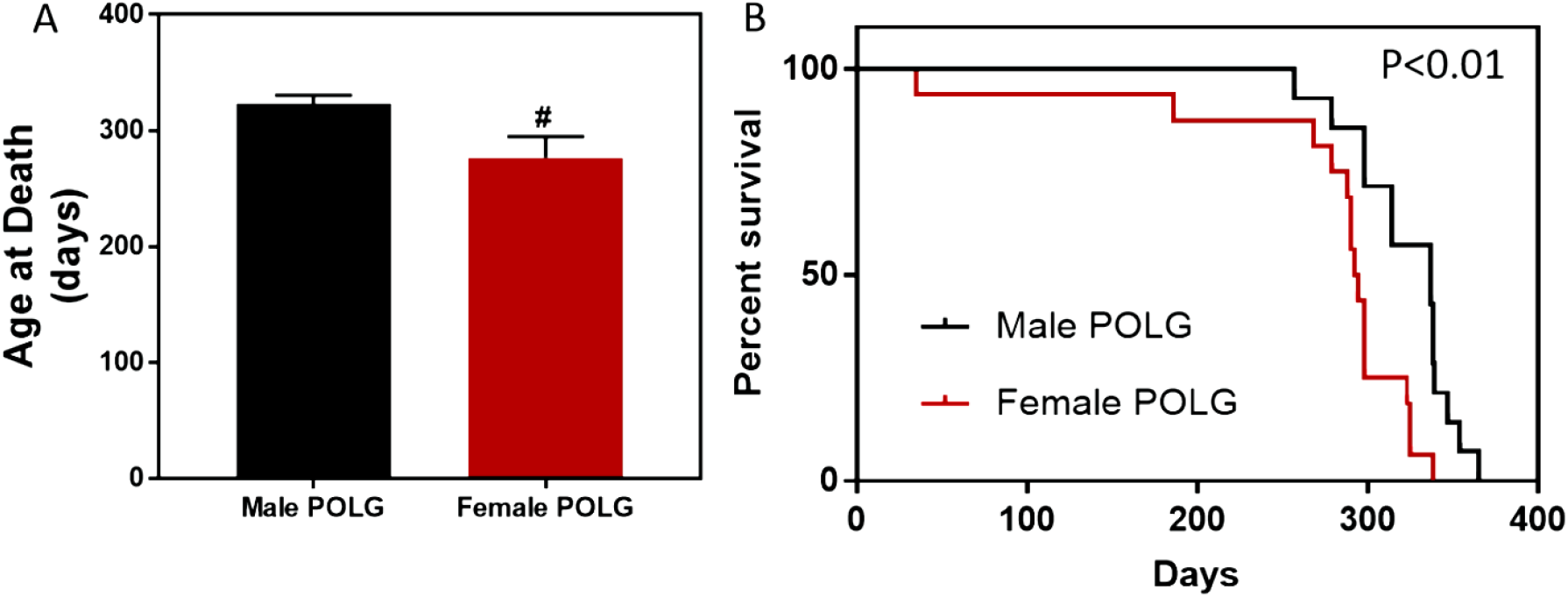
Sex difference in POLG mutant survival. Female POLG mutant mice lived an average of 277 days while male POLG mutants lived an average of 323 days (A, # indicates p<0.05, t test). Survival analysis also showed that female mutants died sooner than males (B, Gehan-Breslow-Wilcoxon p=0.0073, Pantel-Cox p=0.0026, N=14 male and N=16 female).

Given these findings, we sought to assess indices of cardiac aging in male and female POLG mutant and WT control littermates. Echocardiographic imaging was conducted every three months beginning at 6 months of age. Due to differential survival, females were subjected to echo imaging at 6 and 9-10 months of age while males were imaged at 6, 9, and 11-12 months of age with the last echo imaging session being timed with onset of accelerated deterioration of condition. Though no differences were noted in males at 6 months of age, females POLG mutants displayed increased left ventricular diastolic diameter relative to WT controls (Figure 2A). At 9 months of age, both male and female POLG mutants had increased left ventricular diastolic chamber dimensions (Figure 2B). Surprisingly, both male and female POGL mutants displayed apparent improved systolic function as measured by ejection fraction (EF) and fractional shortening (FS) relative to WT littermates (supplemental Table 1). However, it is likely that these changes reflect compensatory mechanisms engaged in the presence of decreased heart rates in the POLG mutants undergoing functional assessment. Also at 9 months of age, female POLG mutants showed increased LV wall thickness relative to controls (Figure 2C). Interestingly, at 11-12 months of age, male POLG mutants displayed LV wall thinning and increased chamber size relative to controls, consistent with a dilated cardiomyopathy phenotype (Figure 2D and 2E). This indicates sex-specific remodeling responses to metabolic dysfunction in the heart.

**Figure 2.**
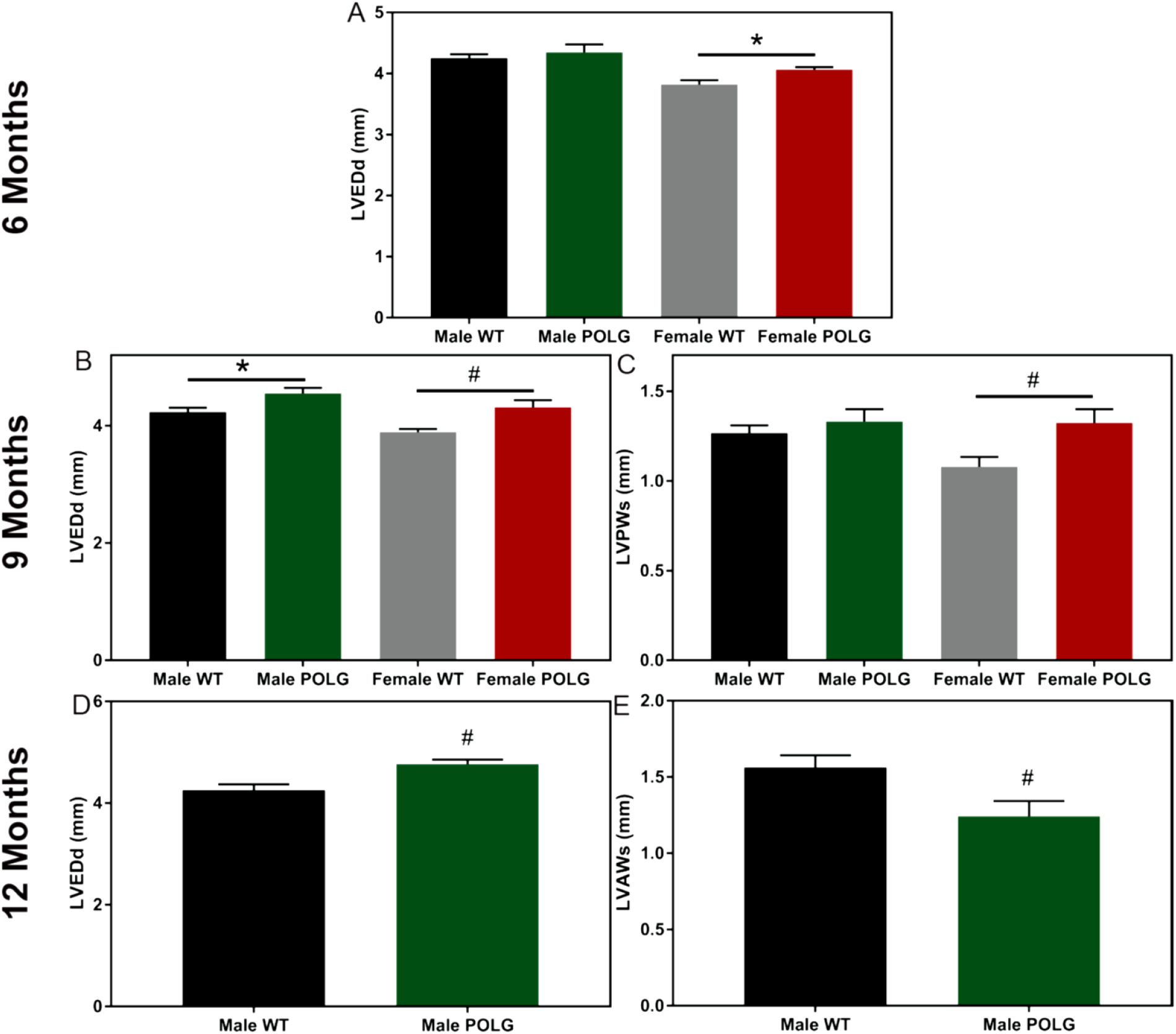
LV chamber remodeling in the POLG mutant. At 6 months of age, female but not male POLG mutant mice had larger LV chambers relative to controls as indicated by diastolic diameter (A). Both male and female POLG mutants displayed larger LV chamber dimensions at 9 months of age (B). Female POLG mutants also displayed LV hypertrophy as indicated by increased posterior wall thickness at 9 months (C). POLG mutant males displayed increased diastolic diameter (D) as well as decreased anterior wall thickness (E). Female mice did not survive to 12 months of age. # indicates p<0.05 and * indicates ANOVA posthoc did not reach significance though t-test was p<0.05 (n=7-10).

Mice were also subjected to invasive hemodynamic measurements of LV and RV function (males 11-12 months of age and females 9-10 months of age). Both male and female POLG mice displayed decreased LV contractility relative to controls as illustrated by decreased Ees and decreased dP/dT max and slowed relaxation as indicated by dP/dT min (Figure 3). In the RVs, despite no change in baseline pressure, the POLG mutants showed decreased dP/dT max and dP/dT min values indicating decreased kinetics of contraction and relaxation, which is consistent with increased Tau values in the POLG (Figure 4). To control for differences in heart rate, dP/dT max and dP/dT min values were divided by heart rate (dP/dT:HR). Still, the POLG mutants displayed decreased contraction and relaxation indices (supplemental Figure 1). Further, LV pressure at dP/dT max (51.6 POLG vs 58.8 WT, p=0.01) and max power (14981 POLG vs 23406 WT, p=0.02) were lower in the POLG mutants, indicating systolic dysfunction.

**Figure 3.**
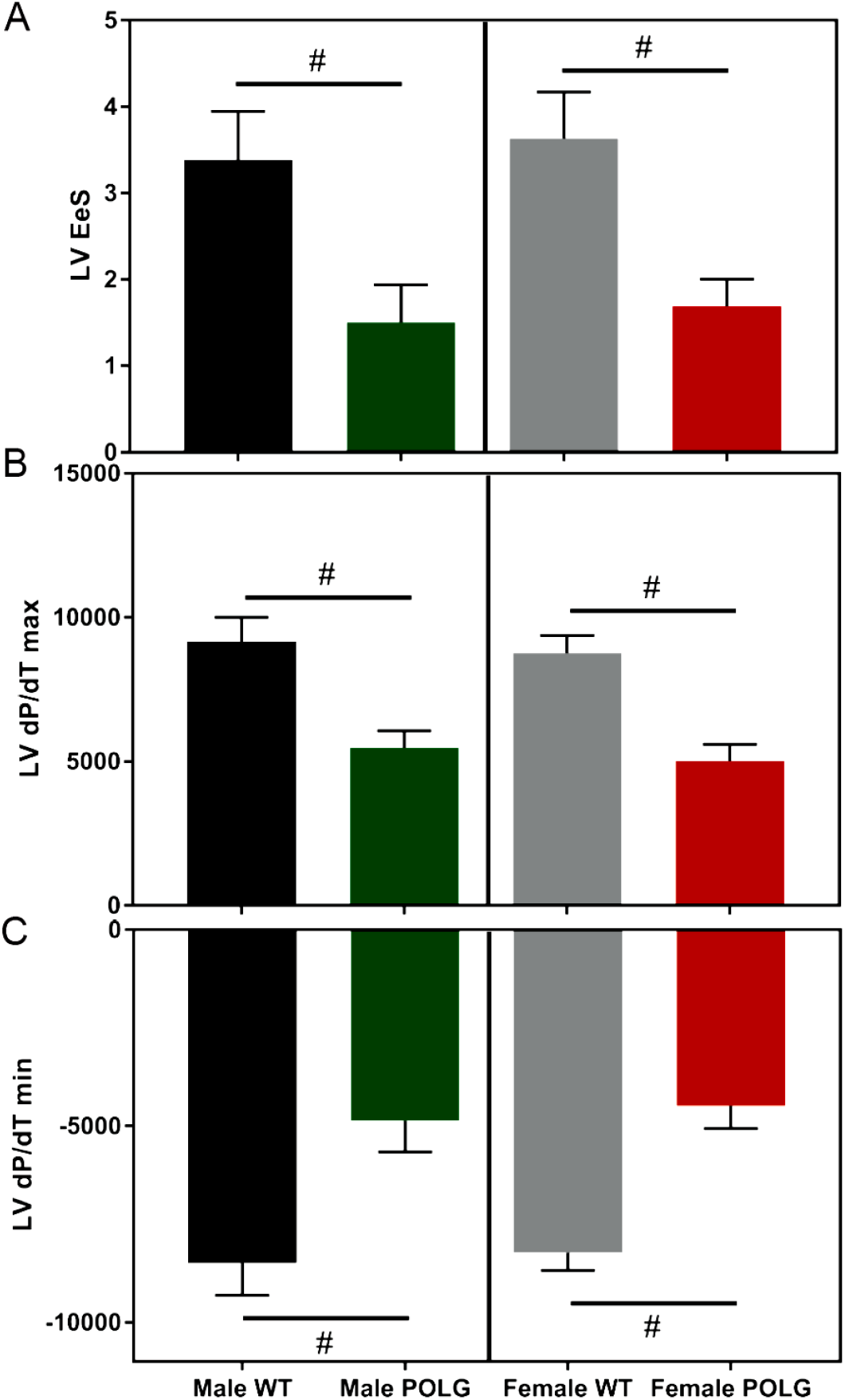
Terminal hemodynamic assessment of LV function. Male and female POLG mutants displayed decreased LV systolic and diastolic function as indicated by decreased Ees (A), dP/dT max (B), and dP/dT min (C). # indicates p<0.05 via t-test vs same sex/age WT littermates. Females are 9-10 months of age and males are 11-12 months of age (n=3 male POLG, n=3 male WT, n=5 female POLG, n=8 female WT).

**Figure 4.**
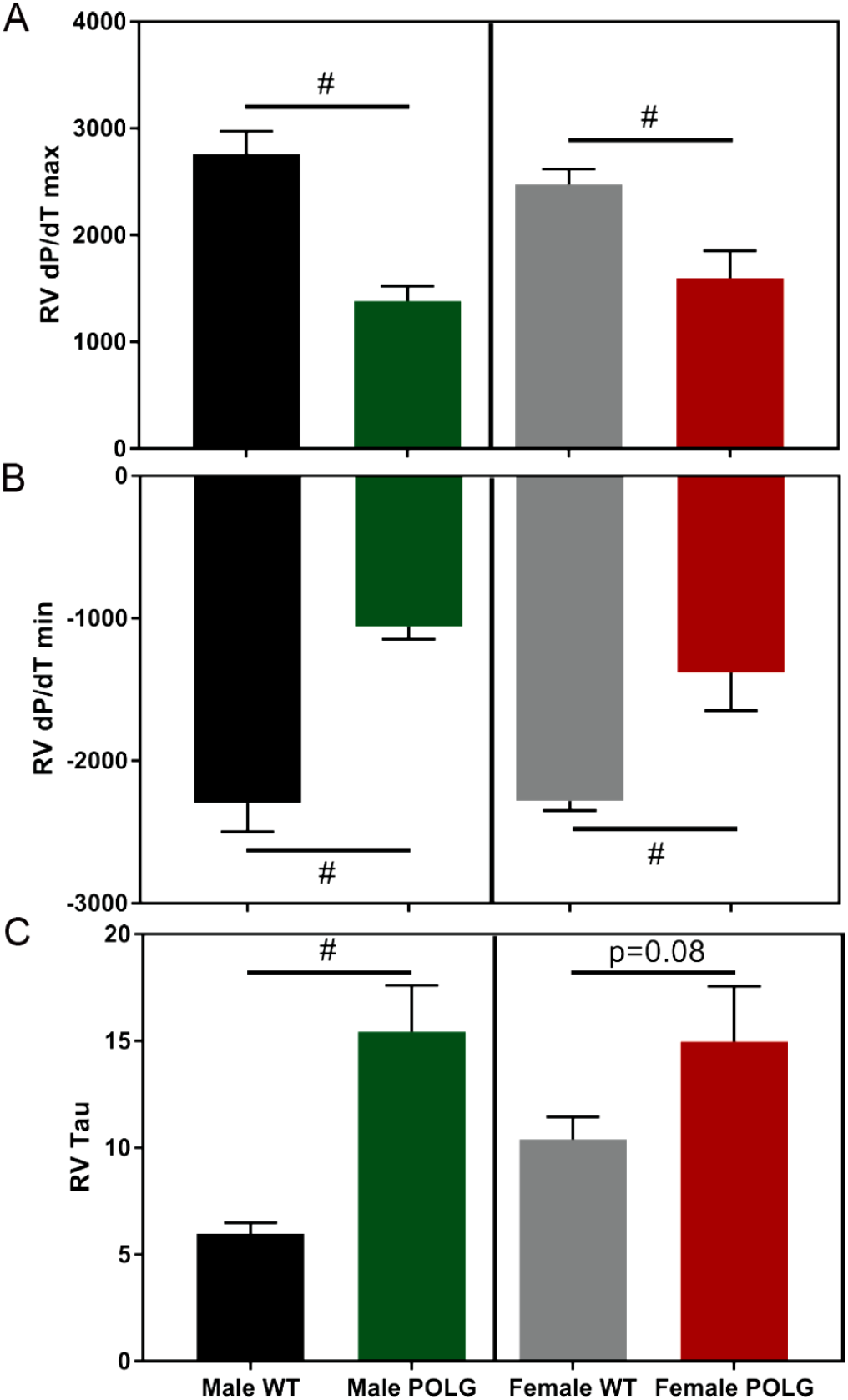
Terminal hemodynamic assessment of RV function. Male and female POLG mutants displayed decreased RV systolic and diastolic function as indicated by decreased dP/dT max (A) and dP/dT min (B). Similarly, Tau, an indicator of time required for relaxation, was increased in the POLG mutant males, though only trended toward an increase in females (D). # indicates p<0.05 via t-test vs same sex/age littermate controls. Females are 9-10 months of age and males 11-12 months of age (n=4 male POLG, n=4 male WT, n=6 female POLG, n=8 female WT).

Consistent with a lack of wall thickening in the male POLG cardiac echos, LV weights normalized to tibia length only indicated LV hypertrophy in female POLG mice (Figure 5A) while RV hypertrophy was noted in POLG mutants from both sexes (Figure 5B). Furthermore, no differences in lung weights (wet weight) suggest that RV remodeling and dysfunction is not secondary to gross pulmonary remodeling or edema (Figure 5C), though liver weights clearly indicate hepatic venous congestion, which is a hallmark of RV failure (Figure 5D, <u>15–17</u>).

**Figure 5.**
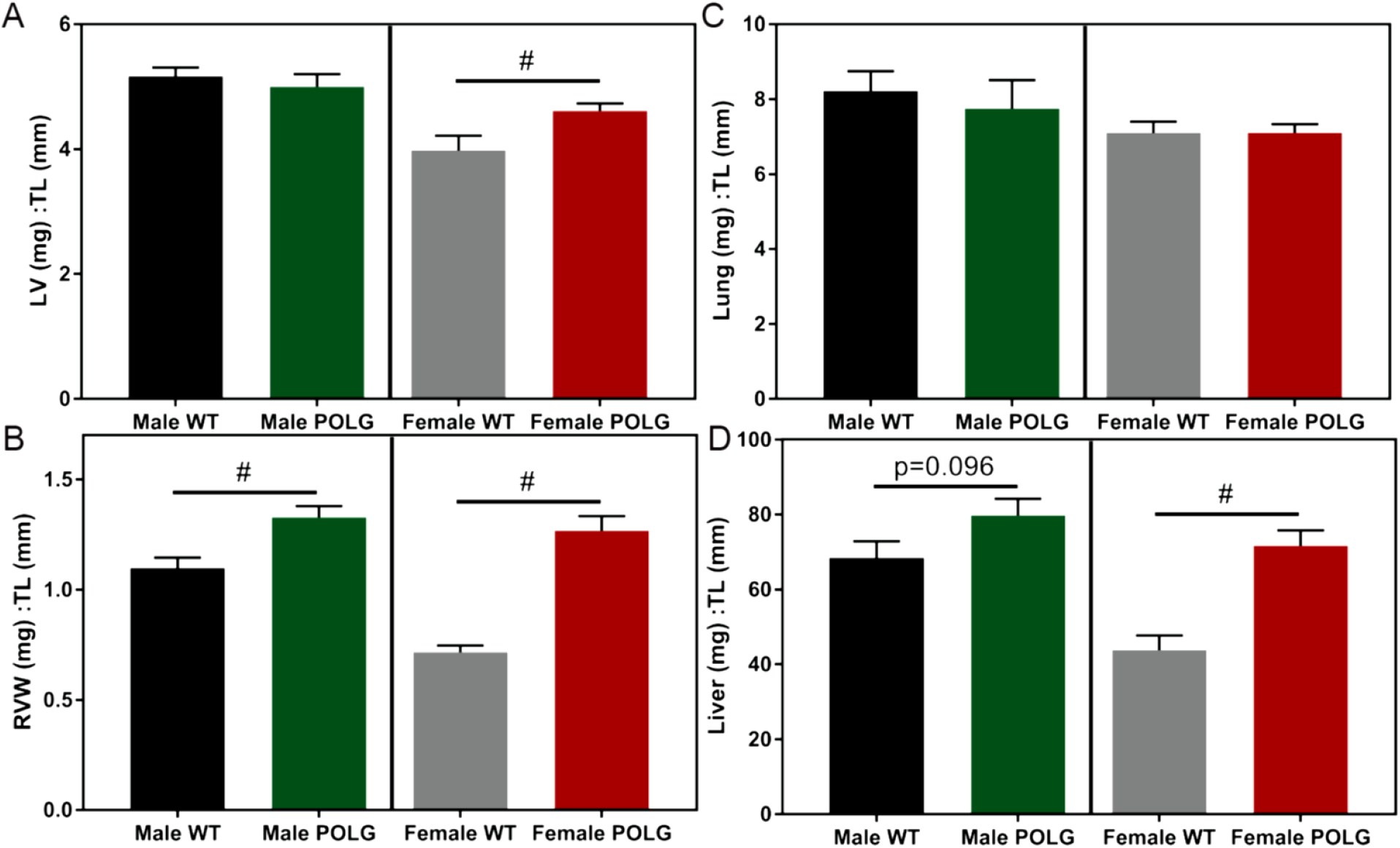
Gravimetric analysis of POLG mutation induced remodeling. Female POLG mutant mice had larger LVs that WT mice while no trend was observed in males (A). Both male and female POLG mutants showed significant RV hypertrophy relative to controls (B) but no indication of pulmonary edema (C). POLG females also showed increased liver weights, indicating hepatic venous congestion, often observed in RV failure. # indicates p<0.05 vs same sex/age WT littermates. Females are 9-10 months of age and males are 11-12 months of age (n=6-7 per group).

To gain insights regarding mechanisms engaged in the presence of severe metabolic dysfunction and associated with RV failure, RNA-seq analysis was conducted on RV tissue in POLG and WT mice (N=4, all females). 455 genes were differentially expressed between WT and POLG RVs (Log2FC>I0.05I, adjusted p<0.05, top 100 displayed in heatmap Figure 6A). Not surprisingly, IPA (Qiagen) analysis of these genes indicated significant alterations in pathways associated with hepatic fibrosis, endocytic pathways, phagosome formation, JAK1/JAK3 signaling, and dilated cardiomyopathy signaling (Figure 6B). Upstream analysis also identified key regulator molecules predicted to be activated or inhibited in the POLG samples relative to controls (Figure 6C and 6D). The statistically strongest prediction was for activation of LPS signaling, a dominant pro-inflammatory molecule. Gene expression changes leading to the prediction of LPS activation are shown in Figure 6E.

**Figure 6.**
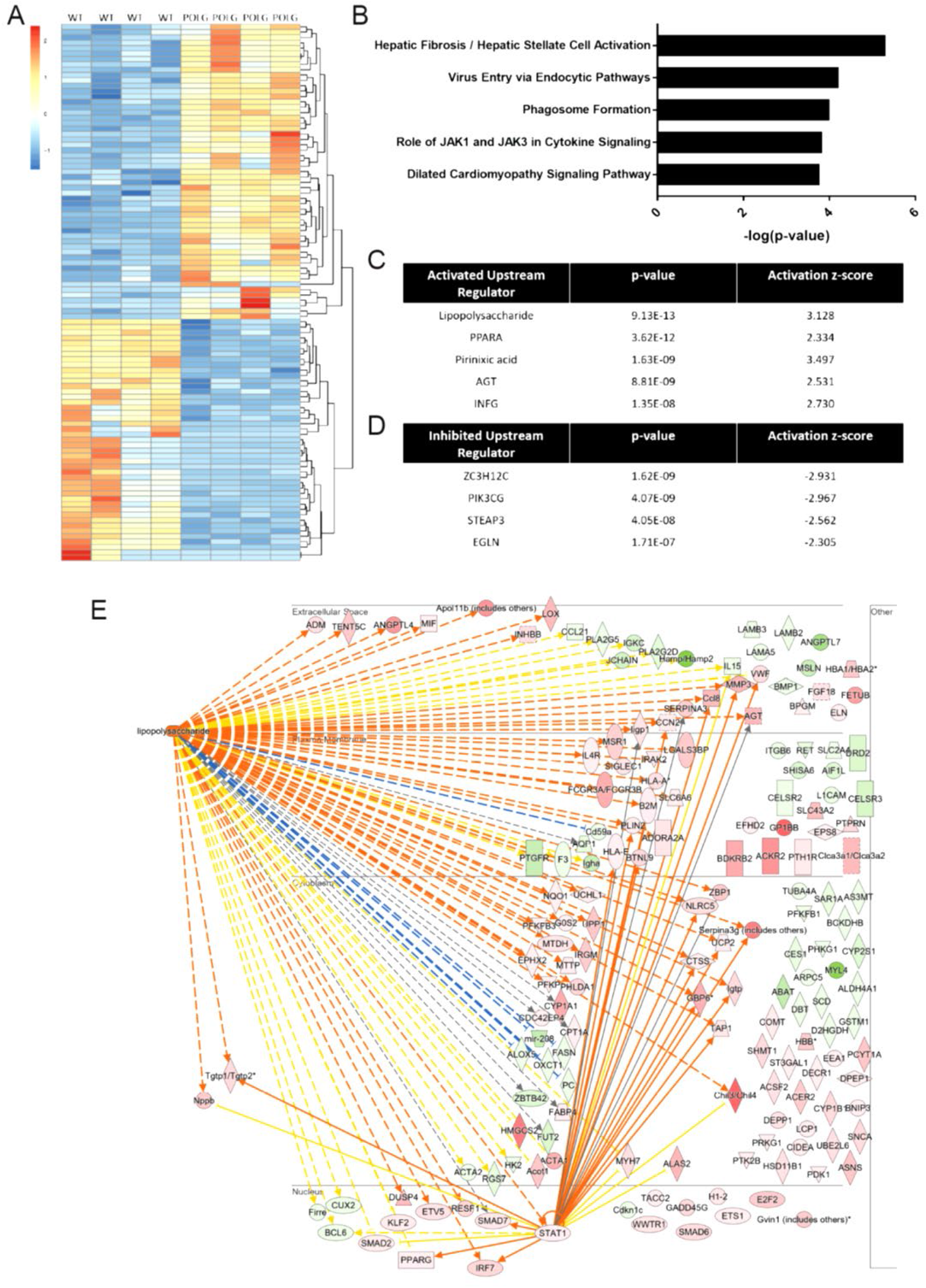
Differential gene expression in the POLG mutant RV. RNA-seq analysis was conducted on RNA isolated from POLG mutant and WT RV tissue. Top 100 of 276 differentially expressed genes (log2FC>I1I, adjusted p<0.05) are depicted in heatmap (A). Core analysis in IPA (Qiagen) was performed on genes showing log2FC>I0.5I (455 differentially expressed genes). Top effected canonical pathways (B) as well as predicted activated upstream regulators (C) and predicted inhibited upstream regulators (D) are shown. Genes leading to the prediction of LPS activation are displayed in (E, gene symbol colors: green indicates decreased expression and red indicates increased expression; line colors: yellow indicates measured expression inconsistent with predicted activity, blue and orange lines indicate measured expression is consistent with predicted activity).

We next sought to compare our transcriptome expression data in the POLG mutant with proteome abundance data in the POLG heart (<u>18</u>) using Gene Set Enrichment Analysis (GSEA, Broad Inst.). A ranked list of genes based on log(2) fold change in the current POLG RNA-seq data (>10,000 genes filtered only by minimum expression) was created and tested against a list of genes encoding significantly upregulated proteins in POLG mutant hearts and again against a list of genes encoding significantly reduced proteins in POLG mutant hearts. Significant enrichment was reached for genes and proteins significantly upregulated by the POLG mutation (Figure 7A, p=0.0058) while genes and proteins with reduced expression were only suggestive of enrichment (Figure 7B, p=0.065). By identifying genes and proteins that clearly move in the same vs opposite directions with POLG mutation, we highlight proteins for which abundance is likely to be regulated by transcriptional vs post-transcriptional mechanisms. Interestingly, proteins that were upregulated consistently with transcription show clear enrichment for platelet related events (Figure 7C, consistent with decompensated heart failure (<u>19</u>), inflammation and tissue damage, reviewed in <u>20</u>) while proteins that are upregulated post-transcriptionally are associated with the mitochondria, particularly protein translation in the mitochondria and ETC assembly (Figure 7C). Proteins that were down regulated in the POLG mutant, regardless of whether the protein change was consistent with the mRNA direction or not, were associated with active mitochondrial function (Figure 7D, e.g. electron transport chain and TCA cycle).

**Figure 7.**
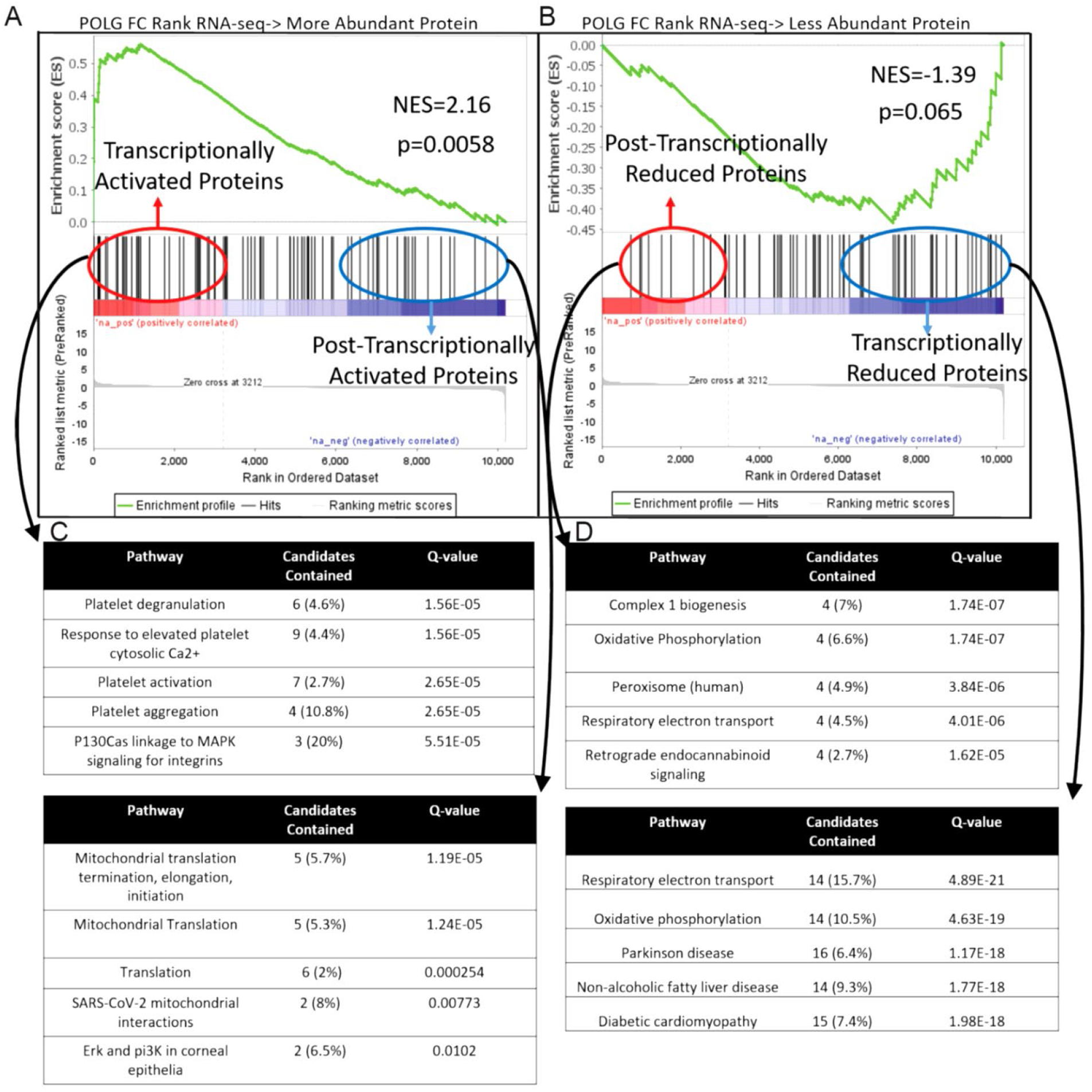
Comparison of POLG effected cardiac Transcriptome vs cardiac Proteasome. GSEA was used to compare the conservation of gene and protein expression changes caused by POLG mutation. Proteins found to be more (A) or less abundant (B) in POLG hearts were converted into gene list and compared against the Log FC ranked RNA-seq dataset. ConcensusPath DB was then used to determine over represented pathways in proteins that were post-transcriptionally vs transcriptionally activated (C) and reduced (D) proteins.

GSEA was also used to determine how closely related the effect of POLG mutation was to accelerated cardiac aging caused in another genetic model (LMNA disruption). Similar to above, a ranked list of genes based on log(2) fold change in the current POLG RNA-seq data (>10,000 genes) was created and tested against gene sets for either LMNA disruption activated genes or LMNA disruption inhibited genes (<u>21</u>). While there was suggestion of enrichment for the genes activated in the LMNA mutant and POLG mutant (Figure 8A, normalized enrichment score (NES) =1.66, p value = 0.065), the enrichment found in inhibited genes was striking (Figure 8B, NES = −1.96, p value = 0.00).

**Figure 8.**
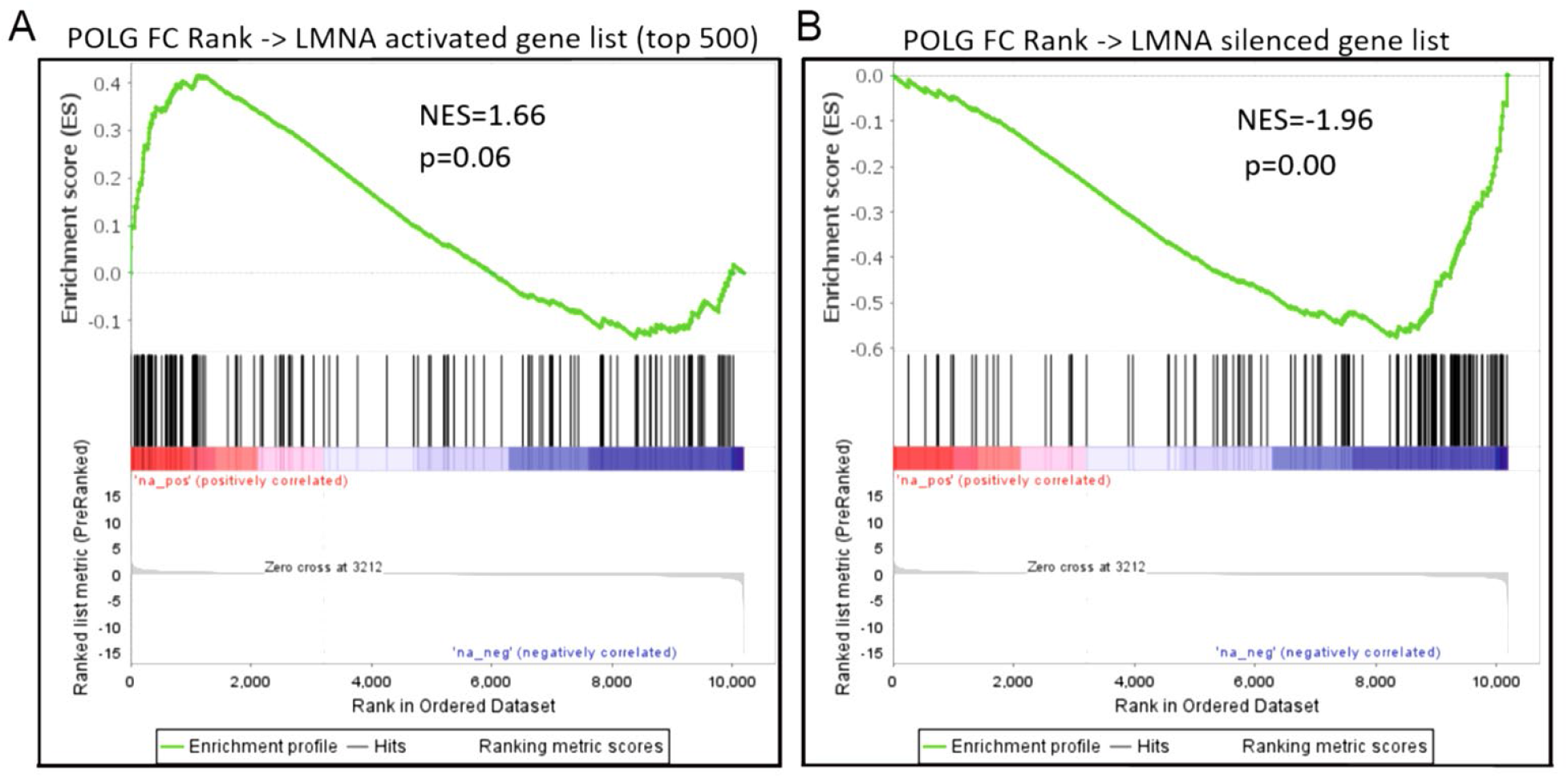
Conservation of gene expression changes in two genetic models of accelerated cardiac aging. GSEA was used to compare the gene expression changes caused by POLG mutation to those caused by cardiomyocyte specific LMNA disruption. While there was suggestion of conservation between the genes activated by both LMNA and POLG mutation (A), the overlap in silenced genes from both models was robust (B).

Given the significant overlap in gene expression between the accelerated aging models, we sought to determine if two promising epigenetic therapeutic approaches might impact significant portions of the POLG mutant gene expression signature (Figure 9). Gene sets for activated or inhibited genes were taken from cardiac HDAC inhibitor (HDACi, ITF2357/Givinostat) and cardiac BET protein inhibitor (BETi, JQ1) treated RNA-seq datasets (<u>22</u> and <u>23</u> respectively). The same POLG log(2) FC ranked gene list as generated for Figures 7 and 8 was compared to these gene sets in GSEA. Both HDACi inhibited and activated genes were significantly enriched in the POLG dataset (HDACi inhibited : NES = 1.39, nominal p value = 0.037; HDACi activated : NES = −1.57, nominal p value = 0.015). The BETi dataset showed marginal enrichment in the POLG data for JQ1 inhibited genes (NES =1.40, nominal p value = 0.09) and also positive enrichment for JQ1 activated genes (i.e. JQ1 activated genes that were also activated by POLG mutation, NES = 1.53, nominal p value = 0.037). This suggests that HDAC inhibitor treatments may provide more benefit in aging associated cardiac pathologies via their ability to rescue the expression of pathologically silenced genes.

**Figure 9.**
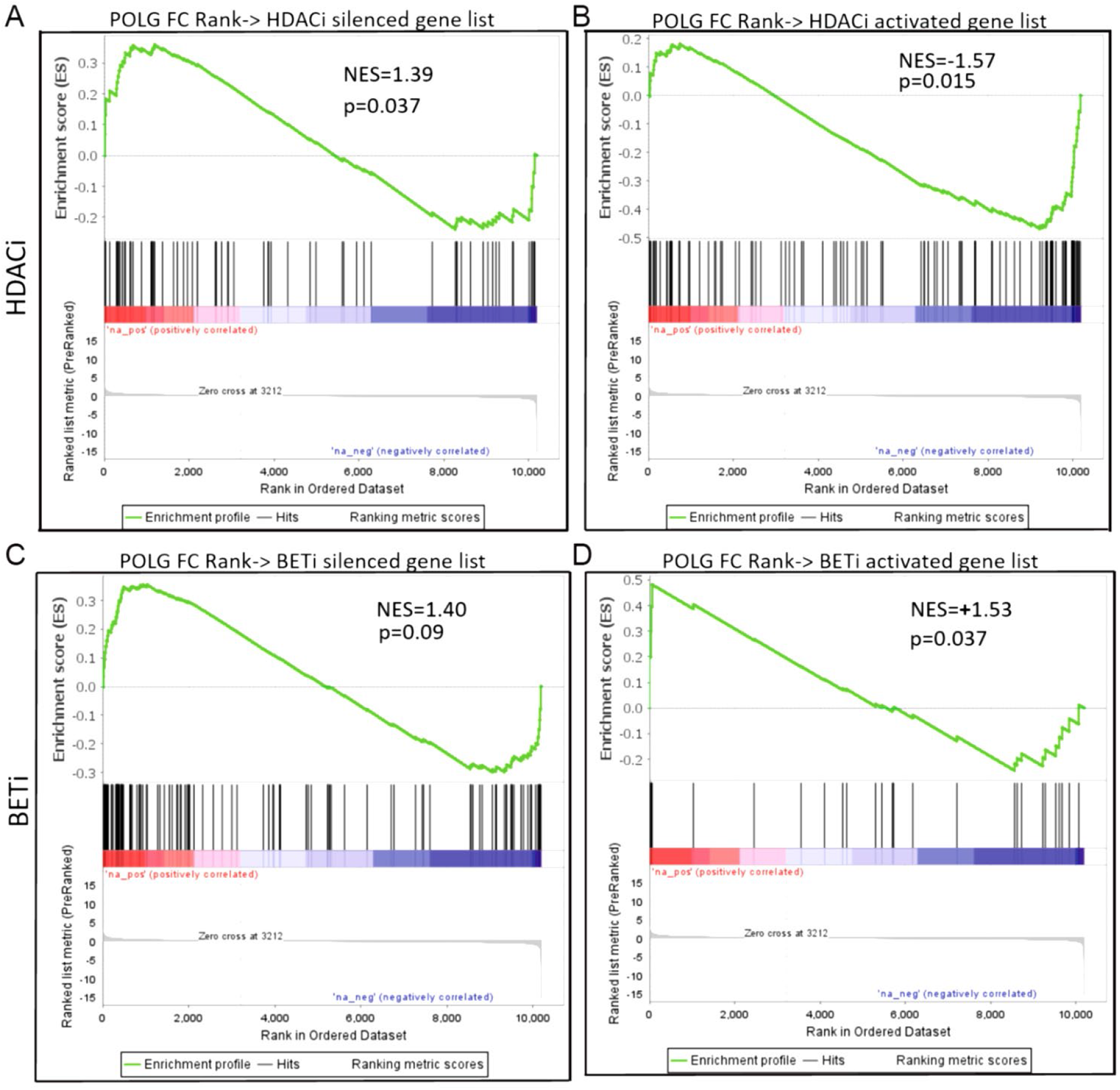
GSEA comparison of POLG effected genes with HDACi or BETi regulated genes. When POLG gene expression changes (fold change ranked gene list) were compared to the genes regulated by HDAC inhibitor treatment in cardiac tissue, significant enrichment was found with HDACi silenced (A) and activated genes (B). No enrichment was found with BET inhibitor silenced genes (C) and positive enrichment was observed for genes activated with BET protein inhibition (D).

## Discussion

Metabolic dysfunction is a component of natural aging. Dysfunction generated during natural aging is typically far milder that what is seen in the POLG mutant (both metabolic and cardiac). We selected this model given the clear cardiac remodeling phenotype and the notion that metabolically driven mechanisms of remodeling associated with aging might be more apparent in this severe model as genome wide epigenetic and transcriptomic studies in natural aging models typically show underwhelming changes.

Several of our findings including *in vivo* survival, cardiac function, and cardiac morphology of POLG mutant mice are not entirely consistent with the interpretations of others who have used this model, which we attribute predominantly to environmental influences alluded to in the introduction (e.g. diet). Additional contributing factors potentially include lack of investigation in female POLG mice, lack of reporting of sex used, method of gravimetrics normalization (i.e. body weight vs tibia length), using echocardiographic calculations of LV mass instead of weights, isoflurane dose used during functional assessment and alternate use of ketamine as anesthetic during imaging. In our hands, mortality was earlier than others have reported and while our IACUC protocol did not support using natural death as an endpoint, euthanized animals were in severely deteriorated condition and unlikely to have survived more than a week past termination. In contrast, others have reported POLG D257A average lifespan at over 400 days (e.g. <u>12</u>). In the referenced study, animals were fed a ~30% fat diet while our animals were fed chow containing only 5.8% fat, potentially suggesting that higher fat content is protective in this model.

Interpretation of the cardiac functional measurements presented are potentially confounded by decreased heart rates observed in the POLG mutants. We attribute this to increased isoflurane sensitivity in the POLG mutants. Future investigation in this model may require cardiac pacing or alternate anesthetic. However, normalization of dp/dt max/min values to heart rate and EeS measurements (less dependent on heart rate) during invasive hemodynamic, as well as elevated liver weights/hepatic congestion support the interpretation of LV and RV dysfunction which is in line with other POLG cardiac literature (e.g. <u>11</u>). Mitochondrial mutation and dysfunction is strongly linked with cardiac dysfunction in the clinic, including for the RV (e.g. <u>24</u>).

Importantly, transcriptome profiling of the POLG RV provides a unique dataset for hypothesis generation regarding gene expression regulation downstream of severe metabolic dysfunction. IPA analysis of differentially expressed genes indicated clear inflammatory gene expression patterns in the POLG mutants as well as significant impacts on fibrosis and cardiomyopathy associated pathways.

Investigation of molecules predicted to be critical up stream regulators strongly support pro-inflammatory signaling activation in the POLG RV. For instance, lipopolysaccharide, AGT (angiotensinogen), INFG were all predicted to be activated and clearly stimulate inflammation (e.g., <u>25–27</u>). Similarly, Zc3h12c, the top predicted inhibited upstream regulator, has been implicated as an inhibitor of NF-kB (<u>28</u>).

Full utility of this RNA-Seq dataset is currently limited by the relative lack of available complementary RV datasets, particularly in accelerated aging models. With this limitation, we opted to compare gene expression changes in the POLG RV with currently available proteomic and transcriptomic data generated from LV or whole heart tissue in models and treatments of interest.

Comparison of gene expression changes in the POLG mutant with protein abundance changes allowed us to identify proteins most likely to be regulated by post-transcriptional mechanisms. Importantly, we identified a number of proteins critical for mitochondrial protein translation that are likely post-transcriptionally activated as these proteins were more abundant in the POLG mutant but their transcript expression was decreased. In contrast, many mitochondrial proteins necessary for metabolic function (e.g. ETC, TCA cycle) were less abundant in the POLG mutant (irrespective of transcript change). We believe this is a compensatory mechanism in the face of severe metabolic dysfunction where the mitochondria elevates expression of proteins associated with mitochondrial protein expression and complex assembly while decreasing expression of the actual subunits the translation and assembly machinery build into functional enzyme complexes. This pattern is strikingly similar to the mitochondrial unfolded protein response (UPR^mt^) where mitochondrial translation- and protein folding- loads are reduced (reviewed in <u>29</u>), though GSEA, IPA, and ConsensusPathDB analyses failed to associate the POLG signature with UPR^mt^. It is likely that this UPR^mt^-like pattern is alternatively activated in the POLG RV as a previous report indicated the main transcriptional activator of UPR^mt^, CHOP/Ddit3, was not elevated in the POLG heart (<u>14</u>) and it is not increased in our data.

Our RNA-seq dataset also allowed comparison of two models of accelerated cardiac aging (POLG and LMNA mutation). Interestingly, while there was some overlap in the upregulated genes in both models, the overlap in downregulated genes in both models was profound. From a philosophical perspective, this potentially indicates cardiac aging is less about what genes are upregulated but rather, function lost due to the genes that have reduced expression. Along these lines, we observed that HDAC inhibition significantly upregulated the expression of genes whose expression was lost in the POLG mutant, reinforcing the notion of zinc dependent HDAC inhibition as a potentially beneficial therapeutic strategy.

This work highlights the degree to which alternate environmental exposures can modulate the presentation of pathology arising from the same genetic mutation and observed sex differences reiterate the complexities of sex-dependent responses to metabolic insult (<u>30–33</u>). Importantly, we provide a unique dataset to the field, which will allow hypothesis generation for investigation of cardiac RV physiology and function. We also conclude that the POLG mutant may be a useful tool for *in vivo* study of non-canonical UPR^mt^, and demonstrate the importance of gene expression rescue in potential anti-aging therapeutics.

## Methods

### Animals

PolG D257A mutant mice were acquired from the Jackson Laboratory (B6.129S7(Cg)-Polgtm1Prol/J, deposited by Dr. Tomas A Prolla) and housed in the Davis Heart and Lung Research Institute vivarium. Wild type (WT) and homozygous POLG D257A mutants (POLG) were generated by heterozygous breeding. Animals were maintained on a 12:12 light:dark schedule and had free access to water and food (Teklad 7012). All animal care and experimental procedures were in accordance with the Ohio State University IACUC and NIH ethical use of animal guidelines.

### Echocardiography

Cardiac function assessment (echo and PV Loop/hemodynamics) was conducted as in <u>Tanwar et al., 2017 (34)</u>. Echocardiography was performed utilizing the Vevo 3100 (Visual Sonics) ultrasound with a 40 MHz transducer. Under light anesthesia (1-2% isoflurane in O_2_), mice were placed in supine position and imaged to aquire LV and RV function. LV chamber dimensions (end-diastolic dimension at diastole [LVEDd], posterior wall at diastole [LVPWd]), fractional shortening (FS) and ejection fraction (EF) were obtained utilizing m-mode in the peristernal short-axis view of the heart at the level of the papillary muscles.

### Hemodynamics

Mice were anesthetized with isoflurane and intubated via a tracheotomy. They were then transferred to a ventilator (VentElite, Harvard Apparatus) at 135 μL/min/g body and kept at 37 °C via a heating pad. PV loops were then performed to evaluate cardiac function with a 1 Fr PVR-1030 small mammal catheter (Millar) and analyzed through LabChart8 Pro (AD Instruments). An IV consisting of 10% Bovine Albumin Serum (BSA) in 0.9% saline was used to maintain fluids throughout the surgery. In some measurements (ESPVR [Ees]) the inferior vena cava was occluded with a suture string to alter pre-load. RV pressure was obtained after measurements were taken from the LV. LV chamber dimensions obtained by echocardiography were used to calibrate LV hemodynamic measurements.

### Terminal Harvest

Under a deep plane of isoflurane anesthesia, the heart was removed and flushed with heparinized saline. Weights were obtained for whole heart, LV, RV, as well as liver (entire) and lungs (entire left and right). Weights were normalized to tibia length. For hearts intended for LV and RV histology, LV/RV dissection was not conducted, otherwise the RV was removed and snap frozen while the LV apex was removed and snap frozen with the LV remainder (mid through base) being formalin fixed for histological analysis.

### RNA-Seq

Sequencing and differential expression analysis conducted by LC Sciences (Houston, TX). Poly(A) RNA sequencing library was prepared from DNAse treated RNA from flash frozen RV tissue (female WT (4) and POLG (4) at 9-10 months of age) following Illumina’s TruSeq-stranded-mRNA sample preparation protocol. RNA integrity was checked with Agilent Technologies 2100 Bioanalyzer. Poly(A) tail-containing mRNAs were purified using oligo-(dT) magnetic beads with two rounds of purification. After purification, poly(A) RNA was fragmented in divalent cation buffer at elevated temperature. Quality control analysis and quantification of the sequencing library were performed using Agilent Technologies 2100 Bioanalyzer High Sensitivity DNA Chip. Paired-ended sequencing was performed on Illumina’s NovaSeq 6000 sequencing system.

### RNA-seq Analysis

Cutadapt (<u>35</u>) and Perl were used to remove adaptor contamination, low quality and undetermined bases. Then sequence quality was verified using FastQC (http://www.bioinformatics.babraham.ac.uk/projects/fastqc/). HISAT2 (<u>36</u>) was used to map reads to the MM10 genome. The mapped reads of each sample were assembled using StringTie (<u>37</u>). Then, all transcriptomes were merged to reconstruct a comprehensive transcriptome using perl and gffcompare. After the final transcriptome was generated, StringTie and edgeR (<u>38</u>) was used to estimate the expression levels of all transcripts. StringTie was used to perform expression leveling for mRNAs by calculating FPKM. Differentially expressed mRNAs were identified by R package edgeR.

### IPA Analysis

Differentially expressed genes were loaded into IPA (Qiagen) and a core analysis was run for genes with adjusted p<0.05 and absolute log2FC>0.5. The top effected canonical pathways and predicted activated and inhibited upstream regulators are reported. Depiction of genes leading to the prediction of LPS activation was generated using the pathway grow and build tool in IPA (intermediate nodes in the pathway that did not show expression or differential expression in the dataset were removed manually).

### GSEA

To compare the POLG mutation gene expression effect to other datasets, a log2FC ranked list of genes from the POLG dataset was compared to gene lists of significantly altered genes/proteins from the datasets indicated in the text using GSEA’s “GSEAPreranked” function (GSEA version 4.2.1, 39, 40). For Figure 7, proteins found to be clearly correlated with transcriptional regulation or post-transcriptional regulation were analyzed in ConsensusPath DB (41, 42).

## Acknowledgements

We would like to thank Prasanjit Sahoo for technical assistance. MSS was supported by NIH AG056848, AHA 857280 and DHLRI institutional funds.

## Data Availability

RNA-seq datasets will be deposited to the NCBI GEO database.

## Supplemental Information

One supplemental table (echocardiography) and one supplemental figure (HR normalization of LV/RV hemodynamic data) are included.

---

**Supplemental Table 1.**

| Age | 6 months |  |  |  |  |  | 9 months |  |  |  |  |  | 12 months |  |  |
| --- | --- | --- | --- | --- | --- | --- | --- | --- | --- | --- | --- | --- | --- | --- | --- |
| Group/Analysis | Male WT | Male POLG | Female WT | Female POLG | Male p = | Female p = | Male WT | Male POLG | Female WT | Female POLG | Male p = | Female p = | Male WT | Male POLG | Male p = |
| Heart Rate | 464.8 | 479.9 | 456.2 | 436.4 | 0.41 | 0.4 | 519.0 | 439.6 | 445.3 | 396.6 | 0.002* | 0.07 | 511.5 | 342.0 | 0.0001* |
| Stroke Volume | 40.5 | 46.5 | 35.8 | 42.4 | 0.1 | 0.01* | 42.0 | 56.7 | 35.5 | 51.5 | 2.7E-06* | 7.4E-04* | 45.9 | 60.2 | 0.008* |
| Ejection Fraction | 49.8 | 54.1 | 57.6 | 58.2 | 0.12 | 0.88 | 52.8 | 61.7 | 54.3 | 60.6 | 0.01* | 0.02* | 56.8 | 61.4 | 0.33 |
| Fractional Shortening | 25.1 | 27.9 | 30.2 | 30.5 | 0.11 | 0.9 | 27.1 | 33.2 | 27.7 | 32.3 | 0.007* | 0.01* | 29.7 | 33.4 | 0.27 |
| Cardiac Output | 18.2 | 22.3 | 16.4 | 18.5 | 0.06 | 0.14 | 21.9 | 24.7 | 15.7 | 20.2 | 0.04* | 0.007* | 23.3 | 20.2 | 0.12 |

**Supplemental Figure 1.**
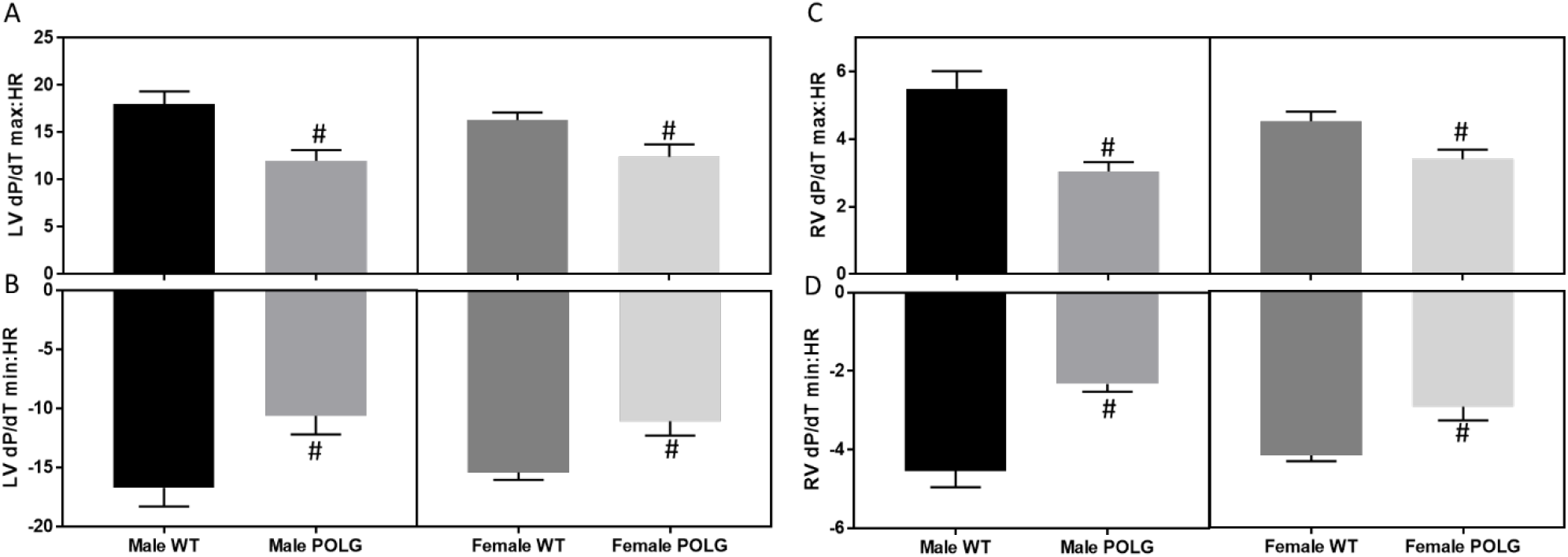
Heart rate normalization of LV and RV function. To normalize potential effects of HR on functional assessment, LV and RV dP/dT max and dP/dT min values were divided by heart rate. LV systolic (A), LV diastolic (B), RV systolic (C), and RV diastolic (D) dysfunction in the POLG mutants remained statistically significant after normalization (relevant to Figure 5 and Figure 6 in the manuscript).

